# Mass extravasation and tubular uptake of red blood cells results in toxic injury to the tubules during kidney ischemia from venous clamping

**DOI:** 10.1101/2022.03.31.486613

**Authors:** Sarah R. McLarnon, Chloe Johnson, Jingping Sun, Qingqing Wei, Gabor Csanyi, Phillip O’Herron, Brendan Marshall, Jennifer C. Sullivan, Amanda Barrett, Paul M. O’Connor

**Affiliations:** Department of Physiology, Medical College of Georgia, Augusta University, Augusta, Georgia; Department of Cell Biology and Physiology, School of Medicine, University of North Carolina, Chapel Hill, North Carolina; Department of Anatomy and Cell Biology, Medical College of Georgia, Augusta University, Augusta, Georgia; Department of Pathology, Medical College of Georgia, Augusta University, Augusta, Georgia; Department of Pharmacology and Toxicology, Augusta University, Augusta, Georgia

**Keywords:** Acute kidney injury, Acute tubular necrosis, Angiophagy, Ischemia-reperfusion, Renal

## Abstract

Vascular congestion is common in ischemic acute kidney injury (AKI) and represents densely packed red blood cells (RBC) in the kidney circulation. In this study we tested the hypothesis that ‘vascular congestion directly promotes tubular injury’.

Studies were performed in male and female Wistar-Kyoto rats. Vascular congestion and tubular injury were examined between renal venous clamping, arterial clamping and venous clamping of blood perfused and blood free kidneys. Vessels were occluded for either 15 or 45 minutes without reperfusion.

We found that venous clamping resulted in greater vascular congestion than arterial clamping, particularly in the outer-medullary region (P<0.001). Venous clamping resulted in significant tubular injury, including cell swelling, tubular degeneration and luminal cast formation following as little as 15 minutes of occlusion. Tubular injury was significantly less following arterial clamping (P<0.001). Numerous red droplets were observed within tubular cells which were most prominent following venous clamping. Electron microscopy and immunohistochemistry identified these as derived from RBCs and indicated that RBCs from congested renal capillaries were extravasated and phagocytosed by tubular cells. CD235a staining confirmed tubular uptake and secretion of RBCs. Cast formation and tubular swelling were absent from blood free kidneys following venous clamping (P<0.001).

Our data demonstrate that congestion of the kidney results in the rapid, mass extravasation and uptake of RBCs by tubular cells causing toxic injury to the tubules. Tubular toxicity from extravasation of RBCs appears to be a major component of tubular injury in ischemic AKI which has not previously been recognized.

## INTRODUCTION

Vascular congestion, or red cell trapping, represents the formation of densely packed red blood cells (RBC) in the renal circulation^1, 2^. Vascular congestion of the outer-medullary (OM) region of the kidney is a hallmark of ischemic acute kidney injury (AKI) with acute tubular necrosis (ATN), now acute tubular injury, in humans at autopsy^3–8^ and is also present in animal models following ischemia reperfusion (IR) from arterial clamping, including in pigs, dogs, rats and mice^9–12^. Vascular congestion of the renal cortical circulation also occurs during periods of ischemia^13^ however, this rapidly resolves following restoration of kidney perfusion^1, 14^.

Our laboratory has recently reported that vascular congestion is associated with severe tubular injury in the renal outer-medullary region and that this occurs within one hour of reperfusion following arterial clamping^14^. It has been speculated that vascular congestion promotes kidney injury by delaying reperfusion and extending ischemic time to congested areas of the kidney, prolonging the period of tissue hypoxia^15, 16^. Opposing this concept however, there is little evidence of medullary hypoxia in the renal outer-medullary region, or other regions of the kidney, in the initial hours after IR^17, 18^. Rather, greater reductions in filtration and subsequent tubular oxygen demands than that of tissue perfusion, have been found to result in increased or stable tissue oxygen levels during this period^17, 18^. The absence of overt hypoxia in the early reperfusion phase, when severe tubular injury is occurring, brings into question the concept that vascular congestion causes tubular injury by prolonging tissue hypoxia.

The goal of the current study was to determine whether vascular congestion causes tubular injury independent of extending ischemia time. As such we compared tubular injury between two models of ischemia, one with severe peritubular congestion and one without significant peritubular congestion. Injury was compared at two time points, 15 and 45 minutes of ischemia, without reperfusion of the kidneys. To promote vascular congestion during ischemia we utilized renal venous clamping. During renal venous clamping, the arterial vessels of the kidney remain unobstructed. As such, blood can enter the kidney but is unable to exit due to the venous occlusion. We hypothesized that this would result in marked vascular congestion during the clamp or ischemic period. Conversely, during the ischemic phase of arterial clamping, blood cannot enter the kidney and vascular congestion of the peritubular capillaries is limited^14^. Only following reperfusion when blood is again able to enter the kidney, does severe vascular congestion of the renal medulla occur^14^. As such, by comparing tubular injury between arterial and venous clamping without reperfusion, we hoped to be able to examine the effects of vascular congestion on tubular injury independent of extending ischemia time. We hypothesized that vascular congestion during the ischemic phase would cause direct tubular injury.

## METHODS

### Animals

All experiments were conducted in accordance with the National Institutes of Health “Guide for the Care and Use of Laboratory Animals” and were approved and monitored by the Augusta University Institutional Animal Care and Use Committee. Age matched Male and female Wistar Kyoto (WKY) rats from Charles River Laboratories were used in all experiments. Rats were housed in temperature (20–26°C) and humidity (30–70%) controlled, 12:12 hour light-cycled conventional animal quarters. Rats were provided ad libitum access to water and standard 18% protein rodent chow (Envigo Teklad, 2918).

### Warm bilateral ischemia-reperfusion surgery and tissue harvest

Ischemia-reperfusion was performed as previously described^14^. Briefly, animals were anesthetized with ~3% isoflurane and 95% oxygen. Body temperature was maintained at ~37°C for the duration of the surgery by servo-controlled heating table and infrared heat lamp (R40, Satco S4998). The renal pedicles were accessed via left and right dorsal flank incisions and blunt dissection with cotton tip applicators. The renal artery was then separated from the renal vein for each kidney using curved forceps. For each animal, the renal artery for one kidney and renal vein for the other kidney were clamped with Schwartz Micro Serrefines (Fine Science Tools #18052-03, Foster City, CA) for either 15 or 45 minutes. The kidney (left or right) which received artery or vein occlusion was rotated for each animal. At the end of the clamp period, the kidneys were excised prior to removal of the vascular clamp. The kidneys were fixed (VIP-Fixative, Fisher #23-730-587) for histological analysis.

In some animals, warm bilateral IR was performed using arterial clamping, and animals were allowed to recover for either 0 (no reperfusion), 1, 2, 6, 10 or 24 hours prior to harvesting the kidney for histological analysis, as previously reported^14^. Corresponding injury and congested data is reported in McLarnon et al^14^.

### Blood free perfusion studies

Animals were anesthetized with ~3% isoflurane and 95% oxygen. Body temperature was maintained at ~37°C for the duration of the surgery by servo-controlled heating table and infrared heat lamp. A midline incision was performed, and the abdominal aorta cannulated. The mesenteric artery and vessels of the right kidney were ligated, and a loose tie placed around the abdominal aorta superior to the renal arteries. The renal vein was separated from the renal artery close to its origin to allow clamp placement. The tie around the aorta was then retracted to prevent blood flow to the left kidney and the kidney flushed of blood via retrograde perfusion of the abdominal aorta using warm ~37°C saline at ~120 mmHg using a syringe heater (EGHSL10, New Era Pump Systems, Farmingdale, NY, USA) and pressure gauge. In blood free animals, the renal vein was clamped during the saline perfusion and the abdominal aorta above the kidney was ligated to prevent blood reperfusion. Renal perfusion pressure was maintained at ~120 mmHg for 5 minutes by adjusting syringe pressure before clamping the renal pedicle for the remaining 40 minutes of ischemia. In control blood perfused kidneys, the kidney was flushed with warm saline but allowed to re-perfuse with blood before clamping the renal vein. Again, the renal pedicle was clamped following 5 minutes of perfusion. At the end of the occlusion period, the kidneys were excised prior to removal of the vascular clamp. The kidneys were then decapsulated, bisected and fixed for histological analysis.

### Assessment of vascular congestion

Paraffin-embedded kidney sections were stained with Gomori’s trichrome (Thermo Scientific, Cat. No.87020), according to the manufacturer’s instructions. Congestion of OM vasa recta (VR) and OM plexus capillaries was assessed using a semi-quantitative scoring method as previously described^14^. Briefly, a score of 0-5 was given to reflect the extent of RBC congestion independently in each region. A score of 0 represents conditions in which all vessels appear open (0% congestion) and a score of 5 represents congestion in all vessels visualized (100% congestion). Scale: 0 = 0%, 1 = 20%, 2 = 40%, 3 = 60%, 4 = 80%, and 5 = 100%. All scoring was done by a researcher unaware of the group identifiers of the samples being scored.

### Assessment of tubular injury

Trichrome stained kidney sections were scored for tubular injury (% injured tubular cells) by a pathologist blinded to the hypothesis of the study and sample identifiers. Tubular injury (cell swelling/necrosis) and tubular cast formation were scored in both the cortex and OM. Tubular injury and cast formation is reported as a score of 0-5 with a score of 0 indicating 0-5%, 1, indicating 5-20%, 2, indicating 20-40%, 3, indicating 40-60%, 4, indicating 60-80%, and 5, indicating 80-100% % of all tubules demonstrating that trait.

### Assessment of red blood cell uptake by renal tubules

To quantify RBC uptake into tubular cells, 3 non-overlapping 100X images of trichrome stained sections of each of the renal cortex, outer-stripe of the OM, inner-stripe of the OM and papilla were taken using an BX40 microscope connected to a DP72 camera (Olympus Imaging Corp, Tokyo, Japan) using an oil immersion lens and Cell Sens imaging software (Olympus imaging Corp, Tokyo, Japan). To avoid selection bias, images were taken by a researcher unaware of the hypothesis being tested or the source of the samples. Images were then viewed with Metamorph imaging software (Molecular Devices, LLC, San Jose, CA) and each cell within an image scored to have either 1) no evidence of RBC uptake, 2) some evidence of RBC uptake, 3) moderate RBC uptake, 4) severe uptake (a tubular cell laden with RBCs). Cells were identified by a nucleus. For each animal, the percentage of cells in each region of the kidney demonstrating either moderate or severe RBC uptake was then calculated.

### CD235a staining

Cut, paraffin embedded sections on slides were heated to 60°C for 30 minutes before being deparaffinized in xylene. Slide were placed in xylene solution 3 times for a period of 5 minutes each. Slides were then rehydrated. Slides were first placed, two times in 100% ethanol for 3 minutes, and then for 3 minutes in 95% ethanol and 3 minutes in 70% ethanol, before being placed in distilled water for 5 minutes. Slides were then placed in antigen retrieval solution (IHC-Tek Epitope retrieval solution (IHC world cat#IW-1100-1L)) for 40 minutes in a steamer, before being allowed to cool for 30 minutes. Slides were then rinsed in 0.1% PBST 3 times for 3 minutes before being placed in 3% H_2_O_2_ solution in distilled water for 15 minutes to inactivate endogenous peroxidases. Slides were then again washed 3 times in 0.1% PBST for 5 minutes each time before being placed in a solution containing 10% goat serum in 0.1% PBST for 30 minutes. Slides were then incubated with anti-CD 235a mouse monoclonal antibody (Invitrogen cat# MA5-12484) at 1:200 dilution in 10% goat serum in 0.1% PBST overnight at 4°C in moist incubation chamber. The next day slides were washed 3 times in 0.1% PBST for 5 minutes each time before being incubated with a secondary antibody (Goat anti-mouse IgG-HRP (Santa-Crus cat#sc-2005)) at 1:400 dilution in 10% goat serum in 0.1% PBST for 50 minutes at room temperature. Slides were then washed 3 times in 0.1% PBST for 5 minutes each time before chromogen staining with 3,3’-Diaminobenzidine (DAB) for 5 minutes. Slides were then rinsed 3 times in distilled water before being counterstained with hematoxylin for 1 minute and washed in running tap water. Slides were then placed in 0.2% ammonia water solution for 20 seconds before being washed again in running tap water for 3-4 minutes. Slides were then sequentially dehydrated in 70% ethanol for 3 minutes, 95% ethanol for 3 minutes and 100% ethanol for 3 minutes, two times before being cleared in xylene solution for 5 minutes x 3 washes. Slides were them covered using Cytoseal XYL medium (Thermo Scientific cat#8312-4).

### Transmission Electron Microscopy

Preparation and imaging of kidney tissue was performed by the Augusta University Histology Core facility. Tissue was fixed in 4% paraformaldehyde, 2% glutaraldehyde in 0.1M sodium cacodylate (NaCac) buffer, pH 7.4, postfixed in 2% osmium tetroxide in NaCac, stained en bloc with 2% uranyl acetate, dehydrated with a graded ethanol series, and embedded in Epon-Araldite resin. Thin 75 nanometer sections were cut with a diamond knife on a Leica EM UC6 ultramicrotome (Leica Microsystems, Bannockburn, IL), collected on copper grids, and stained with uranyl acetate and lead citrate. Tissue was observed in a JEM 1400 Flash transmission electron microscope (JEOL USA, Peabody, MA) at 120 kV and imaged with a Gatan 1095 OneView camera (Gatan, Pleasanton, CA).

## RESULTS

### Renal venous clamping results in greater vascular congestion in the cortex than does renal arterial clamping

To investigate the mechanism(s) through which vascular congestion causes tubular injury, we first compared vascular congestion induced by occlusion of the renal artery to congestion induced by occlusion of the renal vein for 15 or 45 minutes, without reperfusion, in male and female WKY rats (**Figure 1**). There were no sex differences in the degree of vascular congestion with either arterial or venous clamping. While moderate vascular congestion was present in the cortical peritubular capillaries and veins following both 15 and 45 minutes of ischemia from either renal arterial and venous clamping, it was only significantly greater following 45 minutes of venous clamping compared to 45 minutes of arterial clamping (**Figure 1.A. & B**).

**Figure 1.**
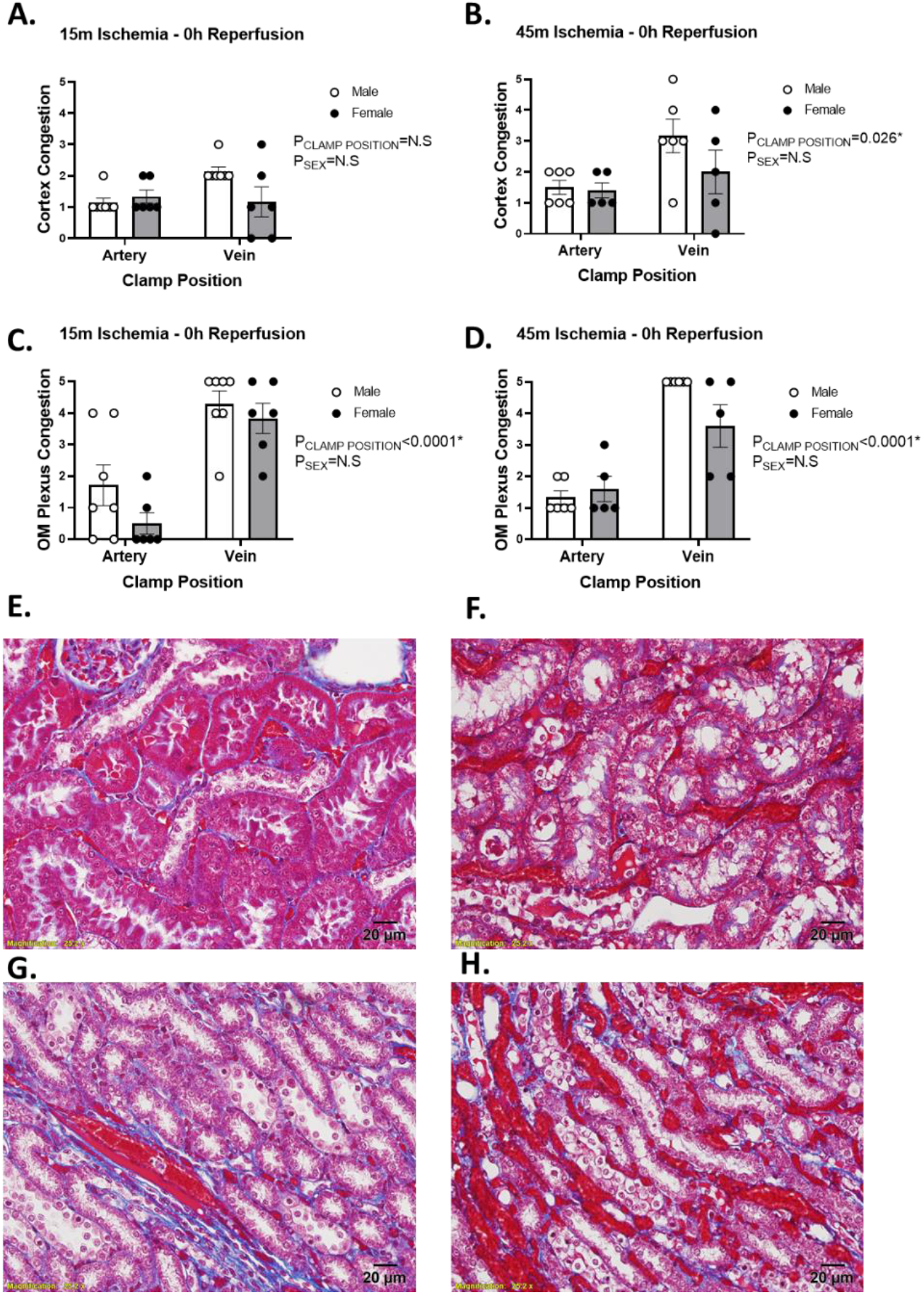
The effect of arterial verses venous clamping on vascular congestion in the cortex and outer-medulla. Vascular congestion scores of the cortex or outer-medullary capillary plexus following 15 (**Panel A and C**, respectively) or 45 minute (**Panel B and D**, respectively) occlusion of the renal artery or renal vein with no reperfusion in male (n=7, white bars) and female (n=6, gray bars) WKY rats. Vascular congestion scores: 0 = 0%, 1 = 20%, 2 = 40%, 3 = 60%, 4 = 80%, 5 = 100%. Values are expressed as mean ± SEM. Two-way ANOVA comparing clamp position and sex, *p<0.05. Note: for each rat, the renal vein was clamped on one kidney and the renal artery was clamped on the other kidney. **Panel E**, Representative images of trichrome stained sections of the cortex following 45-minute occlusion of the renal artery with no reperfusion. There is only moderate congestion of the cortical capillaries. **Panel F**, Representative images of trichrome stained sections of the cortex following 45-minute occlusion of the renal vein with no reperfusion. Cortical capillaries are congested with RBC. **Panel G**, Representative images of trichrome stained sections of the outer medulla (OM) following 45-minute occlusion of the renal artery with no reperfusion. Only the vasa recta bundles appear congested. **Panel H**, Representative images of trichrome stained sections of the outer medulla (OM) following 45-minute occlusion of the renal vein with no reperfusion. Vasa recta bundles and capillary plexus are packed with red blood cells. Images were taken at 40x. Scale bar denotes 20μm.

### Renal venous clamping results in greater vascular congestion in the OM plexus than does renal artery clamping

Renal venous occlusion resulted in significantly greater OM plexus congestion following both 15 (**Figure 1.C, P<0.0001**) and 45 minutes (**Figure 1.D, P<0.0001**) of clamping. There was severe congestion in the OM plexus following 45 minutes of venous clamping, while OM plexus congestion was minimal following arterial clamping. The larger VR vessels were congested following both arterial and venous clamping; however, the VR were more engorged with RBC and VR congestion was more prominent following venous clamping (**Figure 1.G-H**). There were no differences in congestion in the OM between male and female rats (**Figure 1.C & D**).

### Renal venous clamping results in marked tubular injury in the renal OM while OM tubular injury was minimal with arterial clamping

Following as little as 15 minutes of renal venous clamping without reperfusion, there was significant tubular injury in the renal OM, including cell swelling and tubular cast formation. In contrast, few tubules in the OM region demonstrated significant tubular injury following 15 minutes of arterial clamping (**Figure 2.A, P<0.0001**). Tubular injury in the OM was increased following 45 minutes of venous clamping with marked tubular degeneration, indicated by pyknotic nuclei, loss of tubular structure, cellular swelling, and cast formation (**Figure 2.B and F**). In contrast, the percent of injured tubular cells remained low, and tubular morphology looked relatively normal, following 45 minutes of arterial clamping without reperfusion (**Figure 2.B and E**). There were no differences in tubular injury between male and female rats at either time point, however tubular injury tended to be greater in males. Tubular cast formation in the renal OM was exclusive to kidneys in which the renal vein was clamped (**Figure 2.C-D**). There were no sex differences in cast formation in the OM following either 15 or 45 minutes of arterial or venous clamping (**Figure 2.C-D**).

**Figure 2.**
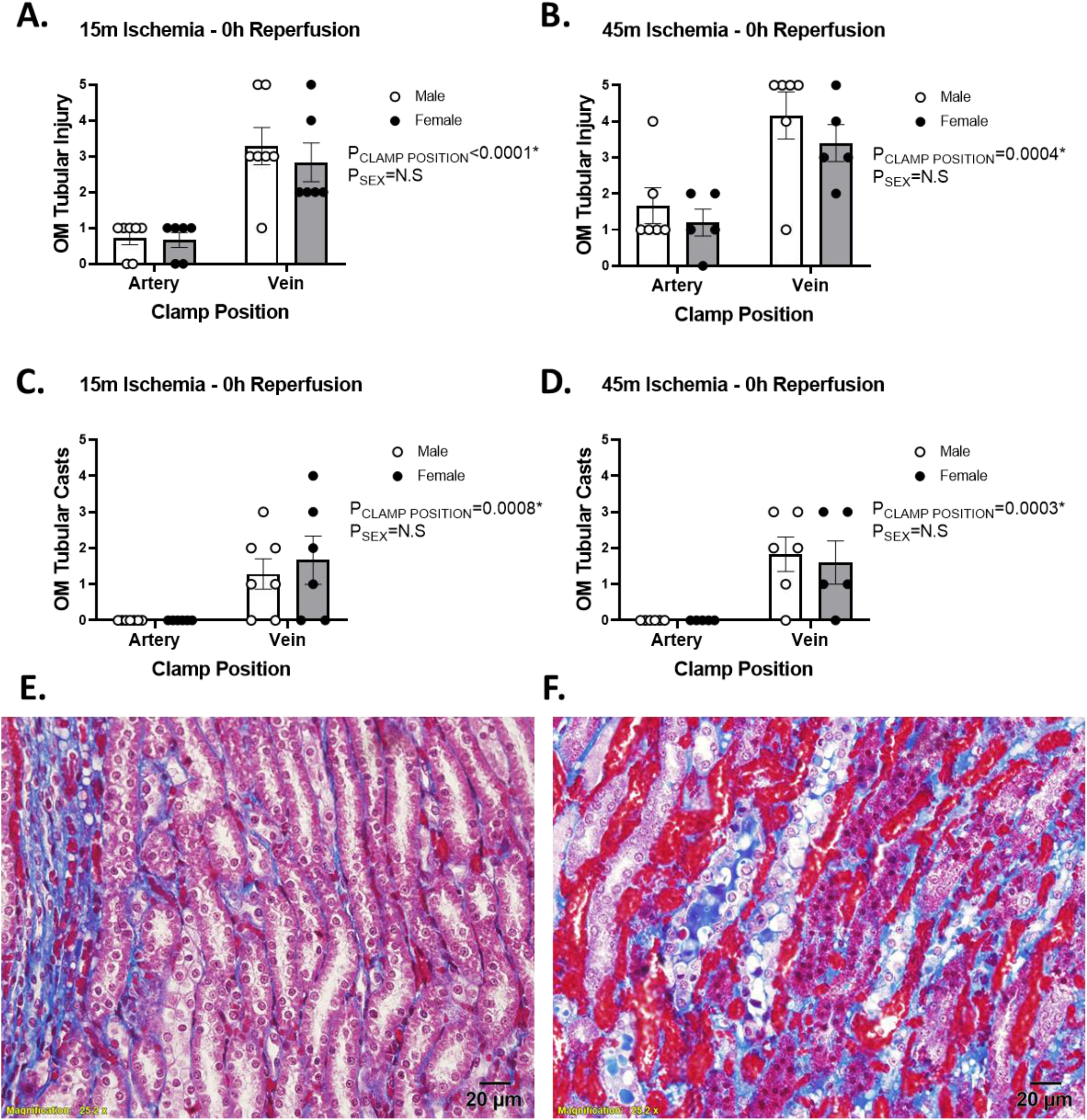
The effect of arterial verses venous clamping on outer-medullary tubular injury. Tubular injury scores of the outer medulla following 15 (**Panel A**) or 45 minutes (**Panel B**), respectively) of occlusion of the renal artery or renal vein with no reperfusion in male (n=7, white circles) and female (n=6, black circles) WKY rats. Tubular cast formation scores of outer medulla following 15 (**Panel C**) or 45 minute (**Panel D**), respectively) occlusion of the renal artery or renal vein with no reperfusion. Tubular injury or cast formation is reported as 0-5, representing the % of tubules demonstrating tubular injury with a score of 0 indicating 0-5%, 1, indicating 5-20%, 2, indicating 20-40%, 3, indicating 40-60%, 4, indicating 60-80%, and 5, indicating 80-100% of all tubules demonstrating that trait. Values are expressed as mean ± SEM. Two-way ANOVA comparing clamp position and sex, *p<0.05. Note: for each rat, the renal vein was clamped on one kidney and the renal artery was clamped on the other kidney. **Panel E** Representative image of tubular morphology in trichrome stained sections of the outer medulla (OM) following 45-minutes of occlusion of the renal artery with no reperfusion. Tubular morphology remains relatively normal following arterial clamping. **Panel F**, Representative image of tubular morphology in trichrome stained sections of the outer medulla (OM) following 45-minutes of occlusion of the renal vein with no reperfusion. Swollen pale cells are common, often with blue or red casts within their lumen. Following 45 minutes of venous clamping, tubular necrosis is also observed, as evidenced by darkened pyknotic nuclei, loss of cellular cytoplasmic area and tubular structure. Images taken at 40X.

### Renal venous clamping results in greater cortical tubular injury compared to renal artery clamping regardless of clamp time

Following 15 minutes of venous or arterial clamping without reperfusion there was significant evidence of tubular injury in the kidney cortex, however, injury was greater with renal venous compared to renal arterial clamping (**Figure 3.A, P=0.003**). In both males and females, all tubules in the renal cortex demonstrated tubular injury following 15 minutes of venous clamping. At this same time point, only approximately 60% of cortical tubules from kidneys with an arterial clamp were injured. Cortical tubular injury was greater in kidneys with arterial clamping following 45 minutes of ischemia when compared to 15 minutes of ischemia, however, the % of injured tubules tended to remain less than that of kidneys with venous clamping (**Figure 3.B, P=0.09**). There were no differences in tubular injury between male and female rats at either time point. Tubular cast formation in the renal cortex was again limited to animals in which the renal vein was clamped (**Figure 3.C-F**). Casts appeared more developed following 45 minutes of venous clamping when compared to 15 minutes of clamping. Following 15 minutes of venous clamping, casts often appeared as discrete droplets within the tubular lumen, rather than the continuous casts filling the entire lumen observed at 45 minutes of clamping (**Figure 3.F**). There were no sex differences in cast formation in the kidney cortex following either 15 or 45 minutes of arterial or venous clamping (**Figure 3.C-D**).

**Figure 3.**
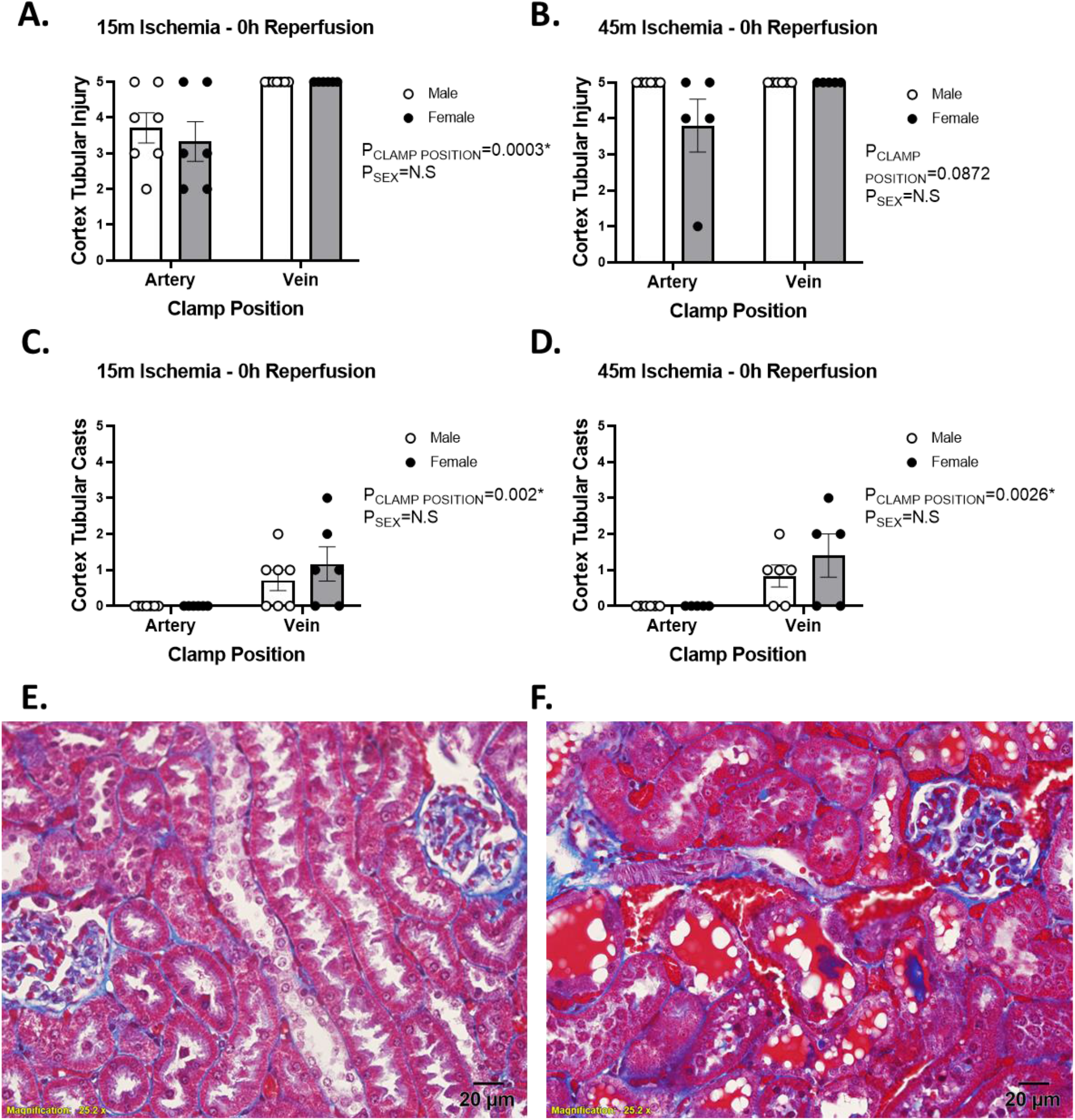
The effect of arterial verses venous clamping on cortical tubular injury. Tubular injury scores of the kidney cortex following 15 (**Panel A**) or 45 minutes (**Panel B**), respectively) of occlusion of the renal artery or renal vein with no reperfusion in male (n=7, white circles) and female (n=6, black circles) WKY rats. Tubular cast formation scores of outer medulla following 15 (**Panel C**) or 45 minute (**Panel D**), respectively) occlusion of the renal artery or renal vein with no reperfusion. Tubular injury or cast formation is reported as 0-5, representing the % of tubules demonstrating tubular injury with a score of 0 indicating 0-5%, 1, indicating 5-20%, 2, indicating 20-40%, 3, indicating 40-60%, 4, indicating 60-80%, and 5, indicating 80-100% of all tubules demonstrating that trait. Values are expressed as mean ± SEM. Two-way ANOVA comparing clamp position and sex, *p<0.05. Note: for each rat, the renal vein was clamped on one kidney and the renal artery was clamped on the other kidney. Images were taken at 40X.

### Tubular cells appeared to take up RBCs and this was enhanced by venous clamping

High power trichrome stained sections revealed the presence of numerous red droplets within tubular cells (**Figure 4.A**). These were present in cortical tubules following both arterial and venous clamping, but were most prominent following venous clamping for 45 minutes (**Figure 4.B & C and F**). These red droplets were also prominent in injured cells or tubules. Some cells appeared so laden with red droplets that they appeared to turn bright red and were almost indistinguishable in color from RBC in surrounding vasculature (**Figure 4.B**). These red droplets were less prominent in the inner-stripe of the OM, being primarily observed in injured tubules following 45 minutes of venous clamping (**Figure 4.D**). Droplets were almost completely absent in tubules of the inner-stripe of the outer-medulla following arterial clamping (**Figure 4.E**). There were no sex differences in the tubular density of these red droplets (**Figure 4.F**).

**Figure 4.**
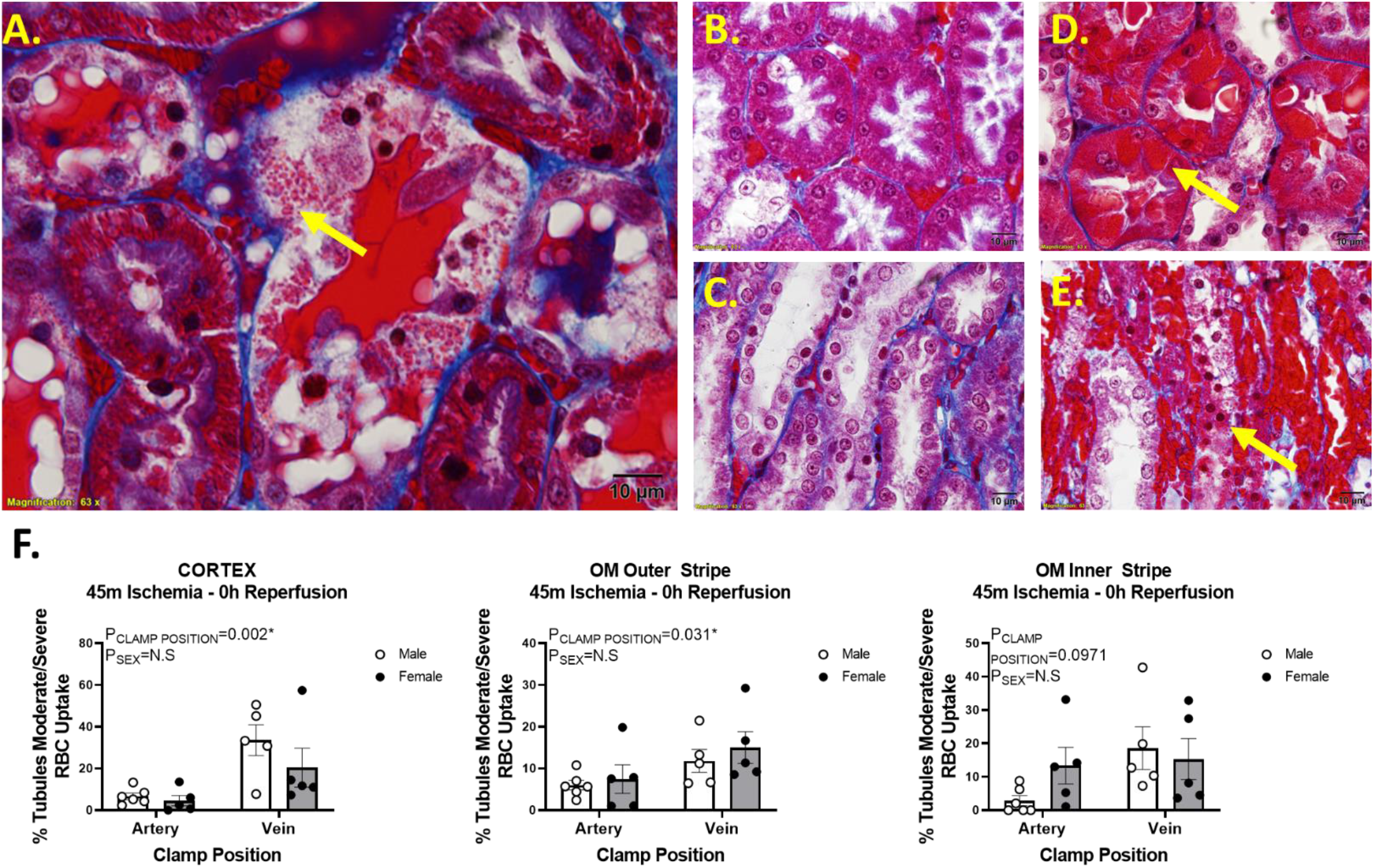
The effect of arterial verses venous clamping on tubular uptake of red blood cells (RBC). **Panel A** Representative image of trichrome stained section of the kidney cortex following 45 minutes of venous occlusion. Red droplets can be observed to have accumulated within the tubular structures, causing them to appear bright red in trichrome stained sections (arrow). **Panel B**, Representative image of trichrome stained section of the cortex following 45-minute occlusion of the renal artery with no reperfusion. Some red droplets are observed in cortical tubules, however the density of these is less than that following venous clamping. **Panel C,** Representative image of trichrome stained section of the kidney outer-medulla following 45 minutes of arterial occlusion. Cells a purple with trichrome staining with little to no evidence of RBC uptake. **Panel D,** Representative image of trichrome stained section of the cortex following 45-minute occlusion of the renal vein with no reperfusion. Tubular cells become deeply red (arrow) and red casts can be observed in the lumen. Note the color change in the cytosol of tubular cells when comparing 45 minutes of arterial or venous clamping as in panels B and D. **Panel E,** Representative image of trichrome stained section of the kidney outer-medulla following 45 minutes of venous occlusion. Cells a purple with trichrome staining with little to no evidence of RBC uptake. Disorganized tubular structure with cells with deep red pigment are common (arrow). All images taken at 100X magnification. **Panel F,** RBC uptake scores of the cortex, outer-stripe of outer-medulla, and inner-stripe of outer-medulla following 45 minute occlusion of the renal artery or renal vein with no reperfusion in male (n=6, white bars) and female (n=5, gray bars) WKY rats. RBC uptake is reported as % tubule cells demonstrating either moderate or severe (RBC laden cells) RBC uptake. Values are expressed as mean ± SEM. Two-way ANOVA comparing clamp position and sex, *p<0.05. Note: for each rat, the renal vein was clamped on one kidney and the renal artery was clamped on the other kidney. All slides were stained at the same time.

### Injury from venous clamping is significantly reduced in the blood free kidney

To determine whether tubular injury from venous clamping required the presence of blood, we compared tubular injury between blood perfused kidneys and saline flushed kidneys that were blood free following 45 minutes of venous clamping without reperfusion. There was significant tubular injury in both the blood free and blood perfused kidney following venous clamping including numerous sloughed epithelial cells which was likely due to the high-pressure perfusion. Cellular swelling and tubular cast formation, however, were almost exclusive to the blood perfused kidney (**Figure 5**). OM tubular cast formation and cell swelling was significantly greater in blood perfused kidneys than saline flushed blood free kidneys following 45 minutes of venous clamping (**Figure 5; P<0.0001**)

**Figure 5.**
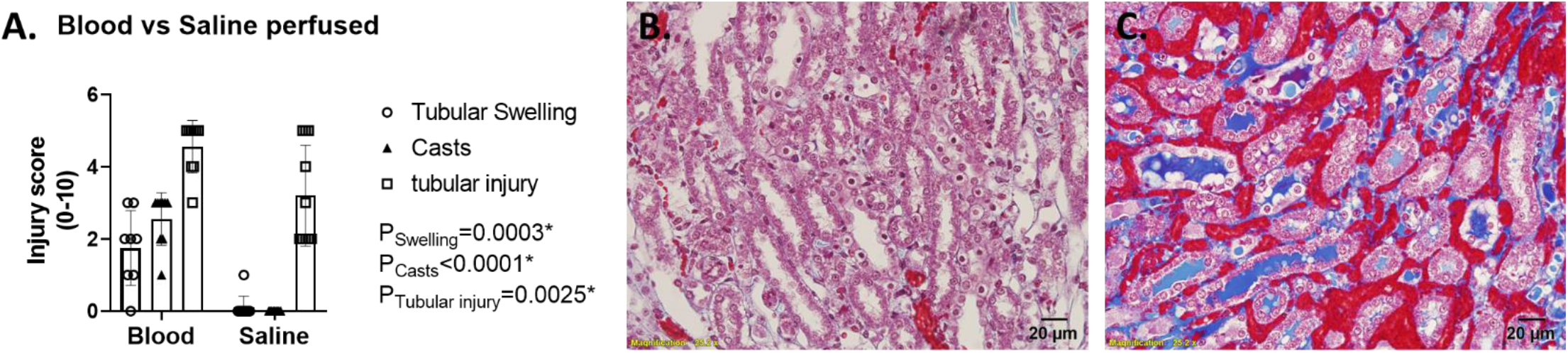
The effect of saline flushing on tubular injury following 45 minutes of venous clamping without reperfusion on OM injury. **Panel A**, Tubular injury scores of the outer-medulla (OM) following 45 minutes of renal venous clamping with no reperfusion in blood and blood free kidneys. As sex differences were not observed, data from male (n=4) and female (n=5) WKY rats were pooled for each panel. Tubular injury is reported as a score of 0-10 with each of cell swelling (open circles), tubular cast formation (closed triangles) and tubular injury (open squares) scored. Values are expressed as mean ± SEM. Statistics represent 2-way ANOVA with Sidaks multiple comparison test, *p<0.05. Representative images of trichrome stained sections of the outer-medulla following 45 minutes of venous occlusion with no-reperfusion in saline flushed ‘blood free; (**Panel B**) and blood perfused (**Panel C**) kidneys are shown. Images taken at 40X magnification. Tubular casts and cell swelling are prominent in the blood perfused kidney but almost completely absent in saline flushed kidneys following venous clamping.

### Electron microscopy imaging of the kidney following 45 minutes of venous clamping demonstrates that RBCs are being taken up by tubular cells and their contents secreted into the tubular lumen

Electron microscopy images from the kidneys of rats following 45 minutes of renal venous clamping without reperfusion identified electron dense components that appeared to be RBCs within tubular cells (**Figure 6.A**). These RBC-like structures within tubular cells often appeared to be in different states of decay, with varying electron density often observed (**Figure 6.B**). Similar electron dense material was often observed within the tubular lumen, suggesting it was being secreted by the tubular cells (**Figure 6.A-C**). Confirming these electron dense structures were RBCs, these structures were completely absence from sections, and the lumen clear of electron dense material, from blood free kidneys following 45 minutes of venous clamping (**Figure 6.D**).

**Figure 6.**
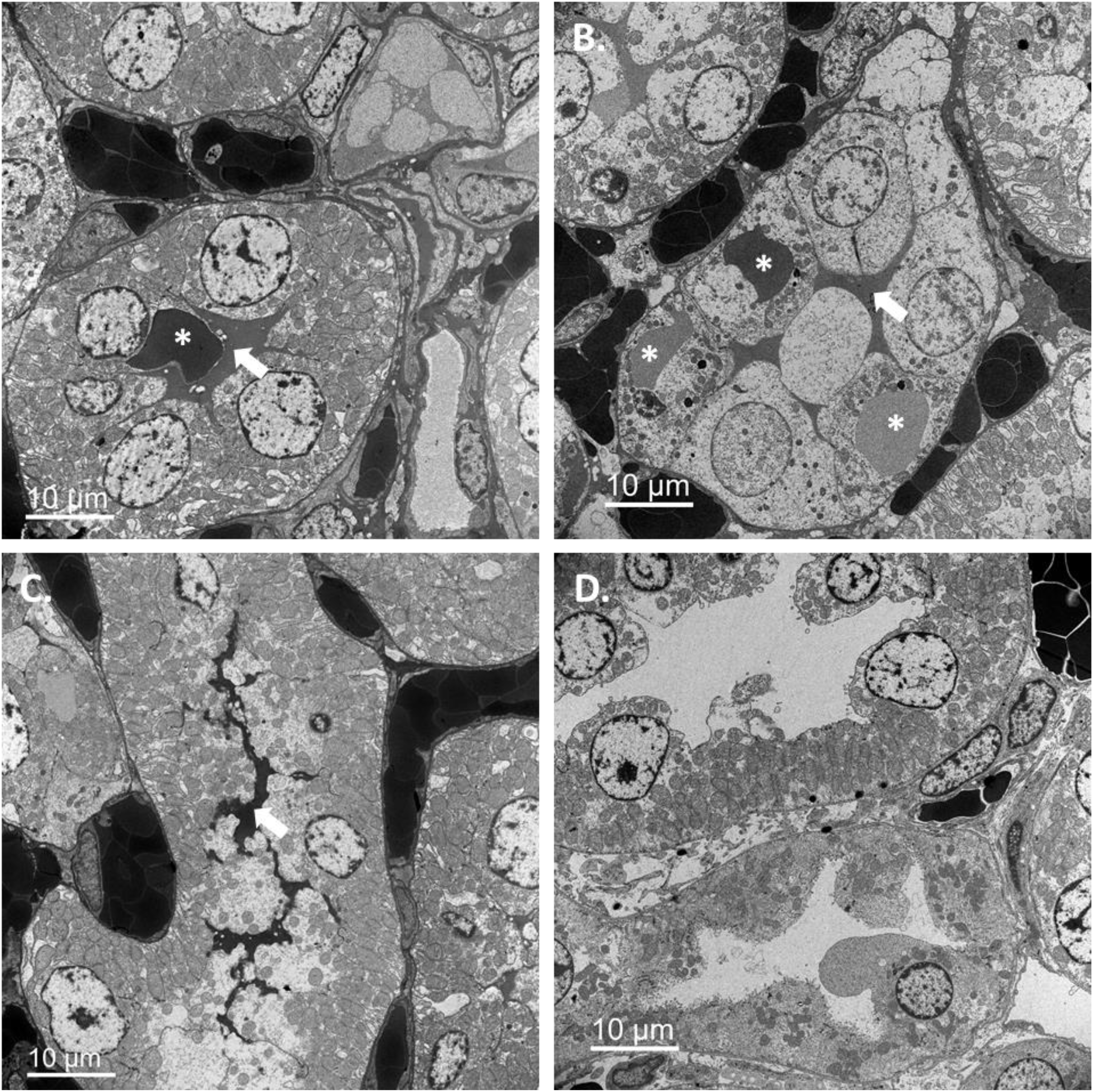
Red blood cells or their components are taken up and secreted by the tubules. **Panels A, B and C** are electron micrographs of the outer-medulla of a rat kidneys 45 minutes after venous clamping. **\*** identify RBCs within tubular cells which appear to be in difference states of decay. **Arrows** point toward RBC like material which appears to be secreted into the tubular lumen. Confirming these electron dense structures are RBCs, these are absent and the lumen open in the blood free kidney following the same 45 minutes of venous clamping (**Panel D**).

### CD235a staining confirmed RBC uptake by tubules

CD235a, or glycophorin A, is a protein which is highly expressed in the cell membrane of erythrocytes. Tubular casts stained strongly positive for CD235a, confirming much of the material within these casts was derived from degraded RBCs (**Figure 7.C-D**). Consistent with uptake of intact RBCs during ischemia following 45 minutes of venous clamping, many tubules had CD235a positive droplets within their cytoplasm (**Figure 7.C and E**). In contrast, CD235a staining was minimal in the renal cortex following ischemia from arterial clamping and positive staining in this group was localized to the vascular compartment (**Figure 7.A**). In the renal OM following ischemia from venous clamping there was diffuse CD235a staining both within tubular segments and in the dilated VR capillaries (**Figure 7.D-F**). Unlike the renal cortex, in the OM CD235a positive droplets within tubules were rare following venous clamping. D235a staining was minimal in the renal OM following 45 minutes of arterial clamping (**Figure 7.B**).

**Figure 7.**
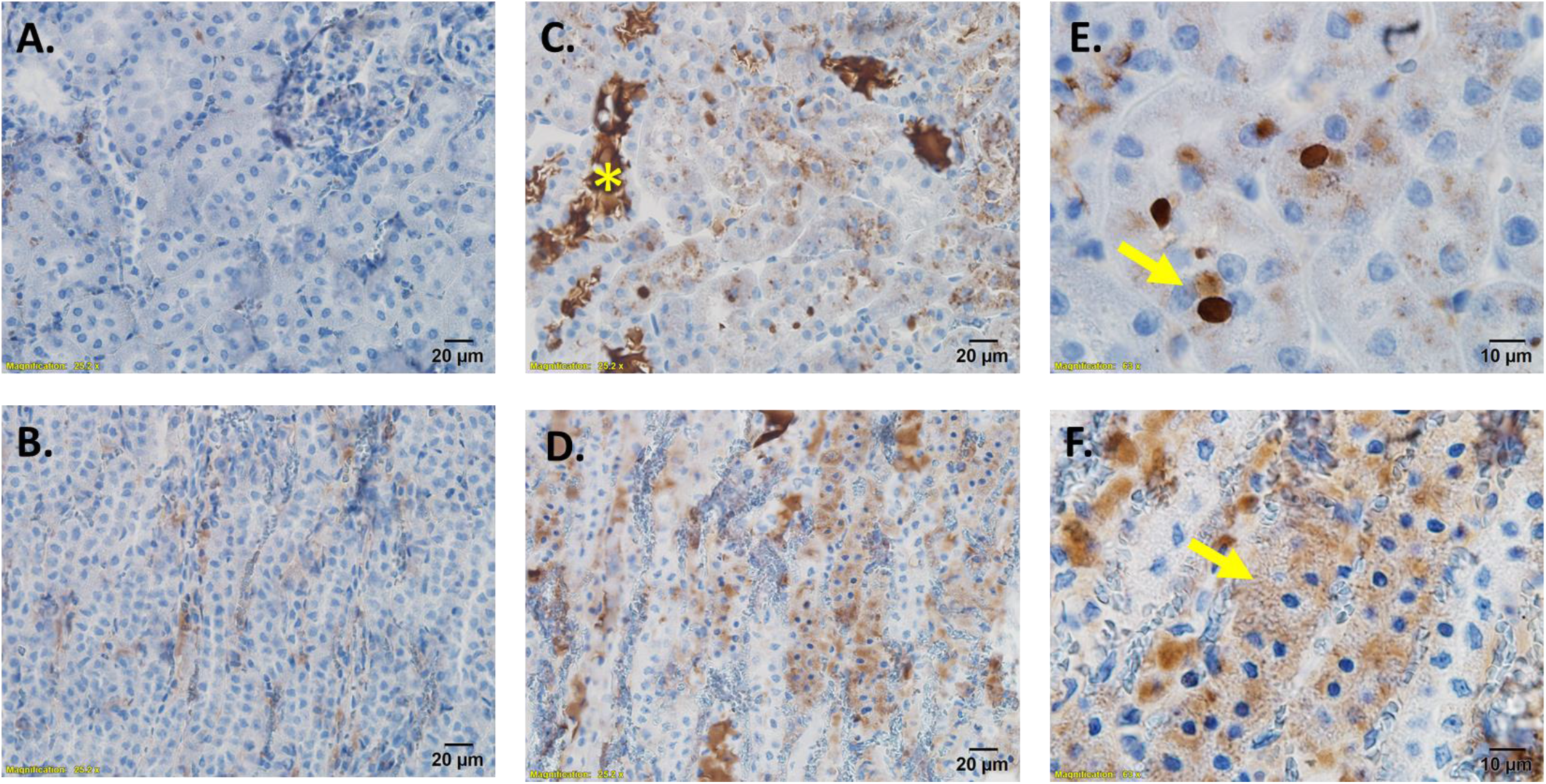
Red blood cell membrane protein CD235a is concentrated in tubular casts and tubular cells following 45 minutes of venous clamping. **Panel A**), CD235a staining in the cortex of a kidney following 45 minutes of ischemia from arterial clamping without reperfusion. Image is 40X magnification. CD235a staining is minimal and localized to within vascular structures. **Panel B**), CD235a staining in the outer-medulla of a kidney following 45 minutes of ischemia from arterial clamping without reperfusion. Image is 40X magnification.CD235a staining is localized to within vascular structures. **Panel C**), CD235a staining in the cortex of a kidney following 45 minutes of ischemia from venous clamping without reperfusion. Image is 40X magnification. Tubular casts stain strongly positive for CD235a. Most tubules also stain positive for CD235a. **Panel D**), CD235a staining in the outer-medulla of a kidney following 45 minutes of ischemia from venous clamping without reperfusion. Image is 40X magnification. Tubular casts again stain strongly positive for CD235a with a diffuse staining pattern observed in many tubules. **Panel E**), Higher magnification (100X) images of cortical tubules following 45 minutes of ischemia from venous clamping without reperfusion. Within cortical tubules CD235a staining is observed within discrete droplets of various sizes. **Panel F**), Higher magnification (100X) images of outer-medullary tubules following 45 minutes of ischemia from venous clamping without reperfusion. Within the outer medulla, CD235a staining within tubules is diffuse and not within discrete droplets.

### Evidence of tubular phagocytosis of RBC and uptake of degraded RBC material between in invaginations of the basolateral membrane

In some images from kidneys following 45 minutes of venous clamping, the basolateral membrane of tubular cells was found to ruffle or closely wrap around what appeared to be intact extravasated RBC in the interstitial space (**Figure 8, A-B**). This suggests phagocytosis of whole RBC may be occurring across the basolateral membrane of tubular cells. Electron dense material from what appeared to be the contents of degraded RBCs within the interstitial compartment was also commonly observed between the folds or invaginations of the basement membrane and appeared to be being secreted through these invaginations into the tubular lumen (**Figure 8, C-D**). In tubular cells with electron dense material filling their lumen, structure similar to the pseudopodia like evaginations reported by Madsen *et al*^19^ in tubular cells phagocytosing RBCs from the apical side could often be observed (**Figure 8.D**).

**Figure 8.**
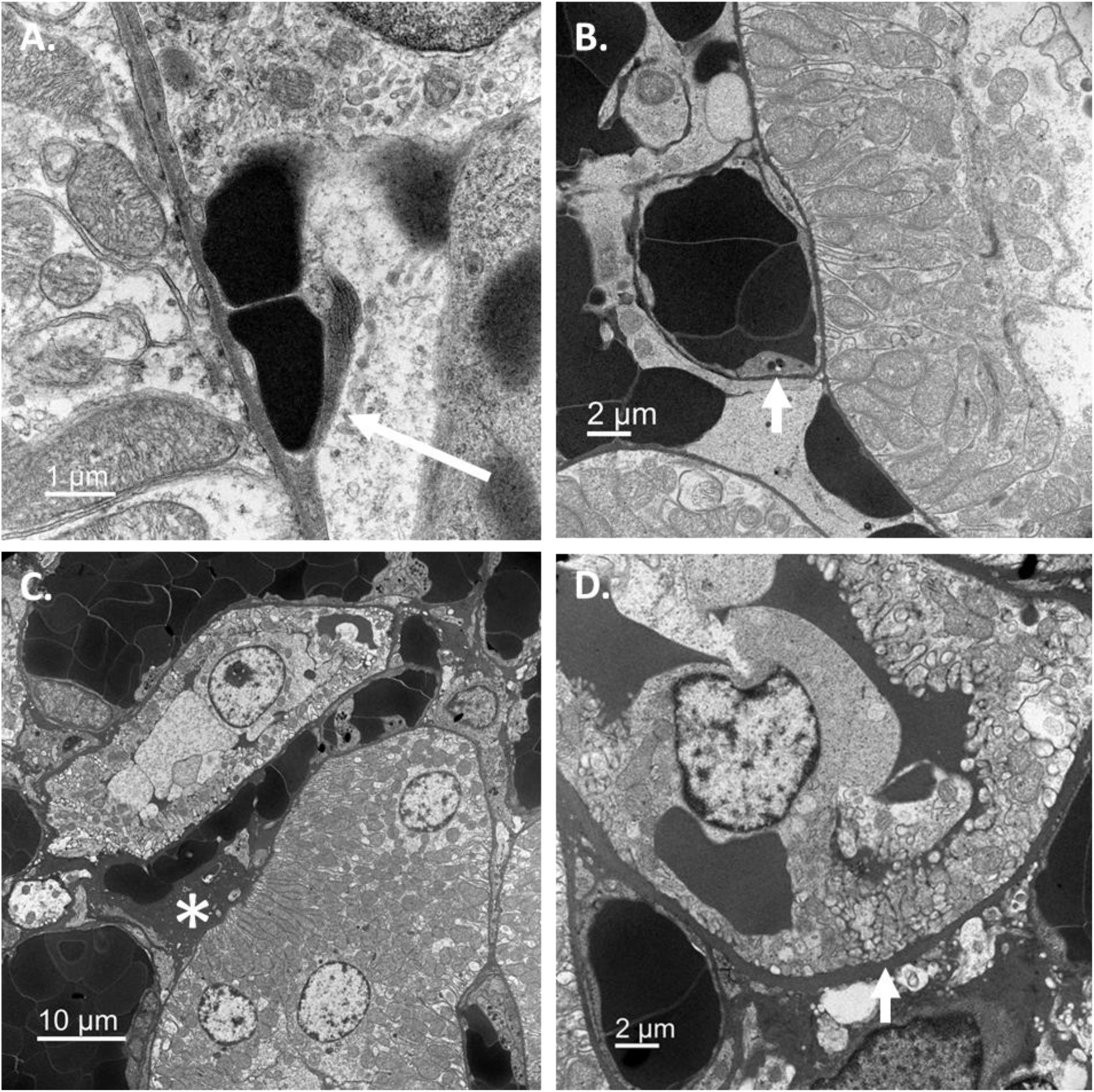
The basolateral membrane of tubular cells was often found to wrap around extravasated RBC and electron dense material could be observed between basolateral membrane invaginations. In electron micrographs of the outer-medulla 45 minutes after clamping the renal vein extravasated RBCs can be observed appearing to be phagocytosed by tubular cells across the basolateral membrane. **Image A**, the black arrow shows the basement membrane of a tubular cell appearing to ruffle and wrap around RBC’s (left image). **Image B**, the yellow arrow (right image) shows the basement membrane of a tubular cell which has completely wrapped around a groups of RBCs. Electron dense material from degraded RBCs was also commonly observed to be taken up between the folds or invaginations of the basement membrane and secreted into the tubular lumen. **Image C**, Electron dense material (*) can be seen outside tubular cells and is continuous with darkly stained basolateral membrane invaginations. **Image D**, a higher magnification view showing electron dense material appearing to traverse between the basolateral and apical membrane between membrane invaginations. In this image pseudopodia like evaginations reported by Madsen *et al*^19^ also appear on the apical side of the cell.

### There is evidence that mass tubular uptake of RBCs also occurs following reperfusion from arterial clamping

Vascular congestion of the kidney cortex and venous circulation occurs during periods of ischemia and marked vascular congestion of the OM occurs following reperfusion from arterial clamping. Further, tubular injury is known to promote later renal vasoconstriction which may result in generalized kidney congestion. To determine if RBC uptake occurs following reperfusion from arterial clamping, we quantified RBC uptake prior to removing the clamp, and at 1, 2, 6, 10 and 24 hours post removal of the clamp, following 45 minutes of warm bilateral ischemia from arterial clamping. At 24 hours’ post-reperfusion from arterial clamping, proteinaceous casts and tubular swelling was prominent. This was associated with histological evidence of RBC uptake like that observed following venous clamping without reperfusion, including many bright red ‘RBC laden’ tubular cells and red droplets within swollen tubular cells (**Figure 9 A-C**). Bright red tubular cells often appeared necrotic and had detached from the basement membrane. Quantification of RBC uptake over time revealed that RBC uptake increased in the more superficial regions of the kidney over the initial 24 hours of the reperfusion period from arterial clamping (**Figure 9.D**). By 24 hours’ post-reperfusion, as many as 40% of tubular cells in the outer-stripe of the OM and 30% of cortical tubule cells displayed significant RBC uptake. RBC uptake in the inner-stripe of the OM was less prominent and was maximal by 2 hours’ post-reperfusion but remained elevated through 24 hours (**Figure 9.D**). A similar pattern was observed in collecting duct cells of the papilla, however, maximal uptake of RBC was not achieved until 6 hours’ post-reperfusion (**Figure 9.D**). Electron microscopy studies of kidney tissue 24 hours’ post-reperfusion from 45 minutes of warm bilateral arterial clamp ischemia demonstrated the presence of electron dense structures within the tubular cytosol of some cells, similar in appearance to dark inclusions found in human kidneys post ischemic AKI (**Figure 9.E-F**)^20^.

**Figure 9.**
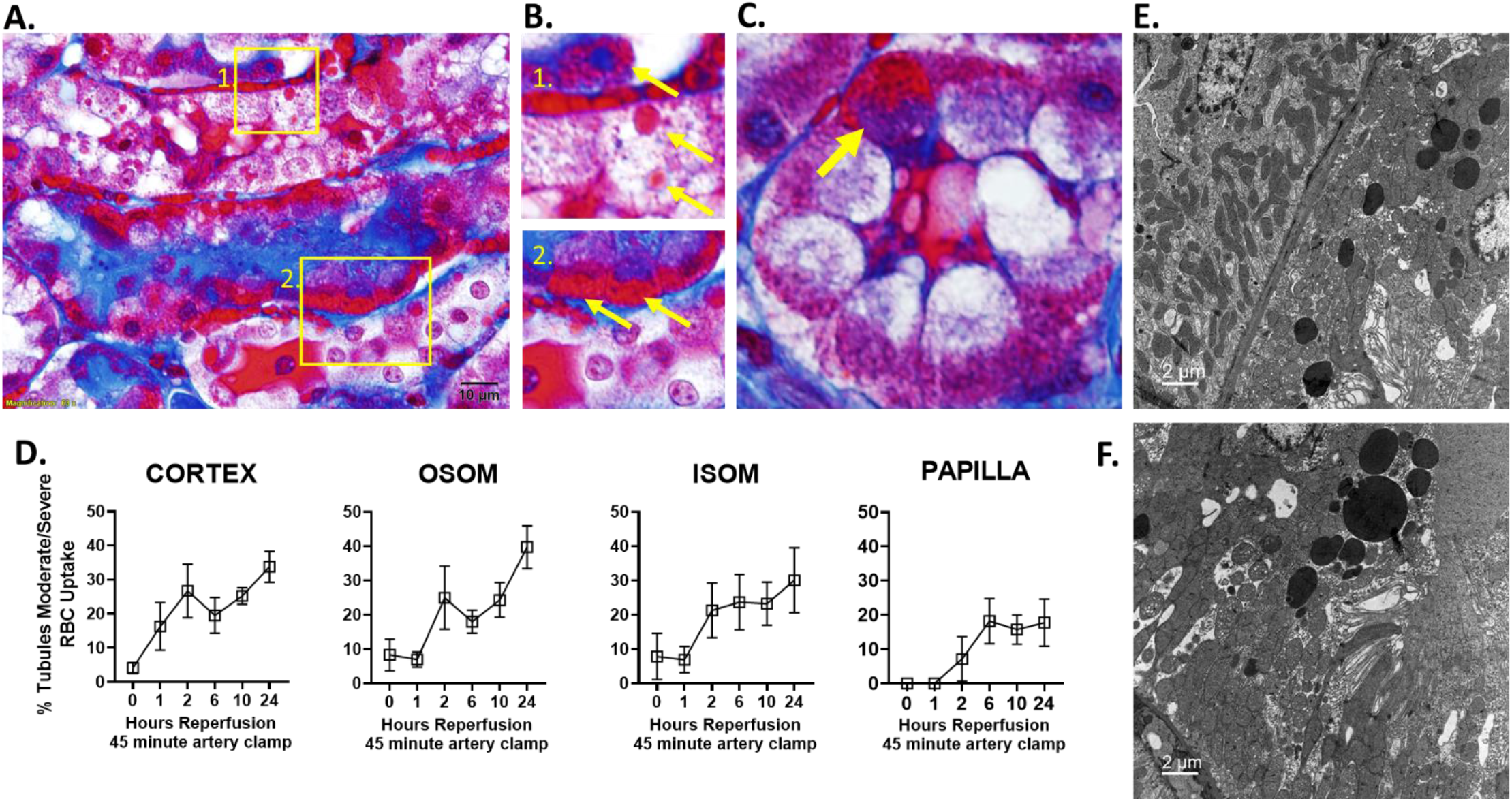
Renal tubular cell uptake of RBCs increases following reperfusion from 45 minutes of warm bilateral arterial clamping. **Panel A** shows a 100X image of the outer-medulla of a rat kidney 24 hours after warm-bilateral IR from arterial clamping. **Panel B** is zoomed images from panel A from the regions depicted in the yellow box labelled (**1.**) and (**2.**) with arrows identifying RBC droplets within tubular cells or RBC laden tubular cells, identified by their bright red trichrome staining. **Panel C** highlights the change in color of a single tubular cell laden with RBC (arrow) compared to other tubular cells in the same structure. In **Panel D**, tubular cells RBC uptake is quantified in the renal cortex, outer-stripe of outer-medulla (OSOM), inner-stripe of outer-medulla (ISOM) and papilla from time=0hr (45 minutes’ ischemia, no reperfusion) through 1, 2, 6, 10 and 24 hours post reperfusion in male WKY rats. Each time point represents quantification of 4-6 animals. RBC uptake is reported as % tubule cells demonstrating either moderate or severe (RBC laden cells) RBC uptake. Values are expressed as mean ± SEM. Two-way ANOVA comparing clamp position and sex, *p<0.05. The percent of tubular cells in all regions demonstrating either moderate or severe RBC uptake increases over the reperfusion period in all regions. In **Panels E-F,** electron microscopy studies of kidney tissue 24 hours’ post-reperfusion from 45 minutes of warm bilateral arterial clamp ischemia demonstrated the presence of electron dense structures within the tubular cytosol.

## DISCUSSION

This study has two major findings: 1) vascular congestion results in rapid extravasation and uptake of RBC by the kidney tubules on mass, in both the cortex and medulla and 2), that this results in toxic injury to the tubular cells. As expected, we found that renal ischemia induced by renal venous clamping resulted in greater vascular congestion than renal arterial clamping, particularly in the OM peritubular capillary networks. Greater vascular congestion in the venous clamp model was associated with significantly greater tubular injury, including cell swelling and luminal cast formation as early as 15 minutes into the clamp period. By the end of the 45-minute ischemic period marked tubular degeneration was observed in the OM along with significant tubular injury and cast formation in the cortex. Demonstrating that this injury was not due to ischemic time alone, despite the same or potentially reduced ischemia time, tubular injury was markedly less following 45 minutes of arterial clamping compared to venous clamping. High power trichrome stained sections revealed the presence of numerous red droplets within tubular cells, which were most prominent following venous clamping. Electron microscopy and immunohistochemical staining for the RBC marker CD235a identified these droplets as RBCs and indicated that RBCs from congested capillaries were being extravasated from the peritubular capillaries and taken up by tubular cells.

While the mechanisms underlying RBC extravasation and uptake remain unclear at present, the basolateral membrane of tubular cells could often be seen to wrap around intact RBCs, suggesting phagocytosis plays a role. Phagocytosis of intact erythrocytes by tubular cells has previously been reported in both rats^19^ and humans^21^ with hematuria, however, to the best of our knowledge, this is the first report of uptake of RBC’s from the capillary network across the basolateral membrane. Electron dense material from what appeared to be degraded RBCs in the interstitial space was also commonly observed to be taken up between the invaginations of the basement membrane of proximal tubular cells and appeared to be being secreted through these invaginations into the tubular lumen. The electron dense pigmented cast material in the lumen could not have been due to glomerular damage as, despite the absence of filtration, it worsened over the ischemic period. Further, following 15 minutes of venous clamping, small bubbles of cast material could be observed which appeared to be being secreted directly from tubular cells. To confirm that tubular injury during ischemia from venous clamping was due to the presence of vascular congestion and RBC uptake, we compared venous clamping between blood and saline perfused, blood free kidneys. Despite the same clamp time, tubular injury was significantly greater, and tubular cast formation limited to the blood perfused kidney.

Our data provide the first evidence that extravasation of RBCs occurs on mass in the kidney within minutes of vascular congestion forming. Grutzendler *et al* recently reported that recanalization of small vessels occluded by emboli occurs by extravasation across the capillary wall in several organs including the kidney^22^. In the kidney, these extravasated emboli are then engulfed by tubular cells over a time frame of many hours^22^. Grutzendler *et al* speculate that this ‘angiophagy’ of emboli, represents and alternative mechanism to hemodynamic forces and the fibrinolytic system to clear obstructions from the microcirculation^22^. Our data indicate that mass extravasation of RBC from congested renal micro vessels begins to occur within minutes of RBC congestion forming. It appears that the mass extravasation and prolonged uptake of RBCs during periods of congestion overwhelms the tubules, leading to toxic injury. In our studies, the tubules of the outer medulla appear to be most vulnerable to RBC toxicity, with many demonstrating severe injury and tubular necrosis despite only modest histological evidence of RBC uptake. Tubules in the cortex appeared to be more resilient, with tubular injury only becoming prominent once cells had become visibly laden with RBCs. We speculate this may be related to the handling of RBCs within the tubular cells. Within cortical tubules, RBCs or RBC material was often observed within discrete droplets, presumably within lysosomes, or between what appeared to be membrane invaginations or channels. In contrast, CD235a staining in the tubules of the OM indicated a diffuse distribution of RBC material within tubular cells, which could result in greater cellular toxicity. The susceptibility of the outer-medullary region to vascular congestion following periods of ischemia^1, 14^, coupled with a lower ability of the tubules in this region to survive toxicity from RBC uptake, provide a novel explanation for the susceptibility of the renal medulla to ischemic injury.

Vascular congestion and tubular RBC toxicity likely explains worsening kidney injury with venous compared to arterial clamping. Clinical reports from partial nephrectomy surgery have suggested that occlusion of the renal vein initiates greater renal damage than occlusion of the renal artery^23^. Studies from rodent models have also shown venous clamping is more detrimental than arterial clamping^3–7^. The pathophysiology underlying worsening renal injury following venous clamping when compared to arterial clamping, however, remains unclear. With venous clamping, blood cannot exit the kidney via the renal vein, however, early in the clamp period, oxygenated blood can still enter the kidney via the artery. This results in packing of the renal vasculature with RBC. This continues until intra-renal pressures become so much that blood flow becomes stagnant. Importantly, as the venous clamped kidney still receives some oxygenated blood early in the clamp period, if ischemia alone was the cause of tubular injury, the arterial clamp would have been expected to result in greater tubular injury than the venous clamp. This, however, is not consistent with the data we obtained. An alternative explanation for greater injury in the venous clamped kidney is that injury is driven by the increased intra-renal pressures. Wei *et al* recently reported that increased proximal tubular pressure during the ischemic phase exacerbates kidney injury during venous clamping^24^. While we cannot exclude a role of increased intra-renal pressures in driving RBC extravasation, in fact we speculate that the increased intravascular pressure is what is driving extravasation of RGB, the absence of tubular cast formation, tubular cell swelling and RBC laden degenerated tubules in saline perfused, venous clamped kidneys, suggests increased pressures alone cannot account for the tubular injury. Rather, our data indicate that RBC mediated tubular toxicity is required for renal injury following venous compared to arterial clamping.

Our data suggest that RBC toxicity is not limited to ischemic AKI from venous clamping. We found evidence that tubular cells also become laden with RBC following reperfusion from arterial clamping. Early in reperfusion from arterial clamping, tubular injury is most prominent in the renal OM. This is where congestion is greatest^14^ and is consistent with our findings in venous clamping indicating RBC congestion results in tubular toxicity. Our data also indicate that tubular RBC toxicity contributes to injury in more superficial nephrons. Cortical tubular RBC uptake appears to increase overtime during reperfusion from arterial clamping. We speculate that RBC uptake in cortical tubules occurs both during the clamp period itself when congestion forms in the cortical vessels, as well as later during reperfusion. This later uptake is likely secondary to injury mediated vasoconstriction that also results in stagnation of flow in areas of the cortical circulation^1, 25, 26^.

RBC uptake and toxicity as a mechanism of injury better accounts for the medullary pattern of injury in rodent ischemia reperfusion models than hypoxic injury. In the arterial-clamp, ischemia-reperfusion model, there is global kidney ischemia during the clamp period. This results in complete cessation of filtration and thus tubular transport. As such, hypoxic tubular injury from the clamp period would be expected to be similar in both cortical and medullary nephron segments. Coupled with the absence of hypoxia during the early reperfusion period^14, 17, 18^, it is difficult to see how the outer-medullary pattern of injury observed in rodent IR models could be attributed to a mismatch in tubular oxygen delivery and oxygen demand for transport leading to hypoxic tubular injury. RBC toxicity provides an explanation for the localization of injury to the outer-medulla, despite global ischemia during the clamp period and the absence of tissue hypoxia during reperfusion^14, 17, 18^ (i.e. OM injury is from tubular toxicity from RBC congestion rather than hypoxic injury).

The contribution of vascular congestion to ischemic AKI in humans remains controversial, however, our findings provide support for RBC toxicity as a key mechanism of injury in human ischemic AKI. Vascular congestion of the OM region is a common finding in ischemic acute kidney injury (AKI) with acute tubular necrosis (ATN) in humans at autopsy from multiple causes ^3–8^. While little is known regarding RBC uptake in human kidneys, RBC like droplets can be observed in injured tubular cells following delayed graft function in trichrome stained sections, suggesting similar processes are occurring^27^. Many of the ultrastructural changes observed in the human kidney following ischemic AKI are also consistent with RBC toxicity. Tubular injury from the luminal phagocytosis of RBCs is remarkably similar to that observed in samples from ischemic AKI, despite the significant absence of glomerular injury in the later. Glomerular bleeding results in tubular phagocytosis of erythrocytes from the tubular lumen which are then broken down within lysosomes^19, 28^. This causes tubular injury, including flattening of the brush border, accumulation of electron dense material within lysosomes and vacuoles, the formation of electron dense pigmented tubular casts in the tubular lumen and cell sloughing^19, 28^. Tubular cells in biopsy specimens from patients with ischemic ATN are also flattened with a loss of brush boarder and marked tubular vacuolization. ‘Dark inclusions’, thought to be degenerate mitochondria^20^, but which we speculate could represent intracellular hemoglobin remaining from the tubular uptake RBCs, have also been reported. Furthermore, Olsen notes, the distal tubules often contain ‘rather dark’ cytoplasm with lumens often filled with ‘chromo-proteinuric’, electron dense, pigmented granular casts, even in cases without myolysis or hemolysis^29, 30^. The ultrastructural commonalities between AKI from the filtration of red blood cells and those from ischemic AKI, including the origin of the electron dense material that forms pigmented tubular casts, can be reconciled if it is assumed tubular RBC uptake was a common element in both forms of injury. The extravasation of plasma and RBCs into the interstitial space by increased capillary pressures is also consistent with reports of edematous interstitial areas with widely separated collagen fibers^20^. Patchy tubular injury with necrotic cells lying adjacent to damaged but intact cells or cells with normal morphology is a well described phenomenon in ischemic AKI. Perhaps most intriguingly in this regard, Dunnill and Jerrome note that ‘the proximity of the peritubular capillaries to the necrotic tubular epithelium could easily be appreciated’^20^. While in their study, they state that this finding is consistent with a back diffusion hypothesis^20^, it would also be consistent with RBC extravasation and nearby tubular cell injury. The extravasation and secretion of RBCs or their components in to the urine during ischemic injury would also explain the ‘heme’ color of ‘muddy brown’ urinary casts which are diagnostic of tubular injury in ischemic AKI^31^. Opposing the concept that vascular congestion and tubular RBC uptake is responsible for injury in ischemic AKI, is the notable absence of vascular congestion many biopsy samples from patients with ATN. There are however potential explanations for this, including; 1) AKI is multifactorial and likely represents a wide range of pathological processes, not all of which may result in vascular congestion and RBC mediated injury, 2) vascular congestion is an early event and is likely to have resolved by the time biopsies are taken, particularly in the more superficial regions of the kidney most often sampled by biopsy, ^1, 2, 14^.

In conclusion, our data, for the first time, demonstrate that RBC congestion of the kidney vasculature results in the mass extravasation and uptake of RBC by tubular cells. The continued uptake of RBCs by most tubular cells during periods prolonged or severe vascular congestion, appears to overwhelm the kidney, resulting in toxic injury to the tubules. The basolateral uptake of RBCs and RBC toxicity appear to be a major component of tubular injury in ischemic AKI which has not previously been recognized. RBC uptake and RBC toxicity as a primary mechanism of tubular injury in ischemic AKI likely explains the greater injury following venous rather than arterial clamping, the relationship between prolonged periods of low renal perfusion and kidney injury^32^ and the failure of diuretics targeting kidney Na^+^ transport and metabolism to improve outcomes in ischemic AKI^2^. Approaches to slow or prevent RBC extravasation and tubular uptake may provide a novel therapeutic pathway to limit kidney injury in ischemic AKI.

## Funding

This work was supported by 1F31DK127683, NIDDK to SM and P01HL134604, NHLBI to PO and NIH NIDDK ISAC GRANT IA2021RC3 to PO.

## Notes

### Competing Interest Statement

The authors have declared no competing interest.

